# VaxKG: Integrating The Vaccine Ontology And VIOLIN For Advanced Vaccine Queries And LLM-Powered Chat Systems

**DOI:** 10.1101/2025.07.15.664450

**Authors:** Feng-Yu (Leo) Yeh, Matthew Asato, Jie Zheng, Yongqun Oliver He

## Abstract

Vaccine research faces challenges in integrating diverse biomedical datasets. While the Vaccine Investigation and Online Information Network (VIOLIN) provides comprehensive vaccine data, implemented in traditional relational models limit complex analysis. Similarly, the Vaccine Ontology (VO) offers standardized semantic frameworks but lacks comprehensive empirical data. This study addresses these limitations by developing the Vaccine Knowledge Graph (VaxKG) that integrates VIOLIN’s dataset with VO’s standardized terminology. Using Neo4j, we transformed 12 core VIOLIN tables into a graph structure enriched with VO concepts. The resulting knowledge graph comprises 28,123 VIOLIN data nodes and 101,282 VO resource nodes, connected by 412,865 relationships. Our comparative analysis of Brucella and Influenza vaccines demonstrates VaxKG’s ability to enable complex semantic queries and reveal insights unavailable from either resource alone. We further demonstrate VaxKG’s utility through VaxChat, a large language model application that leverages the VaxKG as Retrieval-Augmented Generation (RAG) for intuitive vaccine information access.

## Introduction

Vaccination is one of the greatest tools to combat infectious diseases, significantly reducing worldwide mortality rates of various infectious diseases, for example, the effective control of COVID-19 pandemic and the elimination of poliomyelitis in the Americas and the worldwide eradication of smallpox^1^. As we enter the age of big data through the exponential growth of biomedical data available, from genomic profiles to immunological responses, vaccine research has transformed into a much more data-driven field. However, with this wealth of information, researchers face significant challenges in integrating and analyzing diverse datasets.

The Vaccine Investigation and Online Information Network (VIOLIN) database has provided the vaccine research community with a vital platform allowing researchers to easily curate, compare, and analyze vaccine-related research data across various pathogens^2^. Despite the breadth and comprehensive dataset available for vaccine researchers from VIOLIN, the use of traditional relational data models falls short in representing the complex nature of biomedical data, thus limiting the researcher’s ability to perform multi-dimensional analyses that could reveal novel insights into vaccine research. To address this issue, the Vaccine Ontology (VO) has been developed as a structured semantic framework that standardizes vaccine information, enabling more robust data interoperability and facilitating systematic data analysis^3^. VO is a community-based, formal ontology that creates a structured vocabulary while standardizing vaccine terminologies and relationships.

While both VIOLIN and VO offer valuable resources independently, their integration presents an opportunity to overcome the limitations of each system by creating a more comprehensive and interconnected knowledge structure. This research aims to address the challenge of fragmented vaccine data by developing a knowledge graph (KG) that combines the extensive dataset of VIOLIN with the standardized semantic framework of VO. Our work is positioned within the growing field of biomedical knowledge graphs, which have shown success in areas such as COVID-19 research^4^ and genomic data integration^5^.

Similar to the advancement of vaccine research, artificial intelligence (AI) technology has also gained significant growth in research, offering promising solutions to our challenge. KG utilizes graph-based structures to model entities and relationships in a flexible way by organizing information through a network of nodes (entities) and edges (relationships). This approach supports vaccine researchers in enhancing their semantic querying capabilities by traversing multiple complex relationship types and discovering patterns that can reveal novel insights between vaccine entities like components, immune response, genes, or clinical outcomes. Knowledge graphs have proven particularly valuable in biomedical applications where relationship-based insights are crucial. For instance, KG-COVID-19^4^ created a framework to integrate biomedical data for COVID-19 responses, while GenomicKB^5^ utilized knowledge graphs to enable efficient queries across human genomic data.

Our research aims to convert the VIOLIN relational database and incorporate VO into a vaccine KG (VaxKG), a graph database, enriching and unifying the platforms necessary to support efficient vaccine development research. In this study, we will describe the strategy for the conversion of the VIOLIN database into KG, detailing the methodologies for data extraction, transformation, and mapping of domain-specific terms to standardized ontologies. We further demonstrate how this platform helps support data-driven vaccine research through different applications. One application is through semantic queries, allowing users to investigate in-depth insights about vaccine information that may not be available if only using one dataset. Another example is VaxChat, a Large Language Model (LLM) that employs the concept of Retrieval Augmented Generation (RAG)^6^ that uses VaxKG as its knowledge base for the retrieval step in RAG, allowing users to interact with the LLM to retrieve necessary data and generate more focused and accurate answers specifically within the domain of vaccine research. By grounding the LLM’s responses in the structured knowledge of VaxKG, VaxChat overcomes limitations of general-purpose LLMs like hallucination and provides reliable and context-aware information.

## Methods

### Data Sources

The VIOLIN database serves as a comprehensive platform to store vaccine-related information, capturing data from various curated tables. Our study utilizes 12 core VIOLIN tables, including *t_adjuvant, t_gene, t_host, t_gene_engineering, t_host_gene_response, t_host_response, t_pathogen, t_reference, t_vaccine, t_vaccine_detail, t_vaxjo* (for vaccine adjuvants), and *t_vaxvec* (for vaccine vectors). These tables collectively provide detailed data on vaccines, vaccine components, host responses, pathogen targets, and vaccine-specific attributes. Before the integration process, raw data was extracted from the VIOLIN MySQL relational database and converted into CSV format, ensuring consistency for further processing.

To enhance semantic interoperability and enrich the VIOLIN dataset, VO (released on 2025-02-26) is also incorporated. The ontology file (in OWL format) was downloaded^7^ and is used as a reference framework for annotating and integrating the vaccine data.

### VIOLIN Data Import

VaxKG was implemented using the Neo4j Desktop for database management, and the import directory was configured to securely host all CSV and ontology files. Data transformation, necessary for converting relational data into graph nodes and relationships, was achieved through custom Python scripts (found in GitHub repository)^8^ that leveraged the Neo4j Python driver. Each table was transformed into a specific node type (e.g., Vaccine, Gene, Host, Pathogen) with associated properties, and relationships were later defined based on foreign key mappings.

### Ontology Integration Method and Mapping

VO integration was performed using the Neosemantics (n10s)^9^ plugin from Neo4j. The process began with configuring n10s to optimize ontology handling within the graph database, setting parameters for Uniform Resource Identifiers (URI) handling, relationship type management, and RDF type interpretation. The VO ontology file was then imported directly from its public file^7^. This would parse the ontology into RDF triples, represented as `Resource` nodes, which form the basis of the VO vocabulary within the structure of the knowledge graph.

Following ontology import, a comprehensive mapping strategy was implemented to link VIOLIN data nodes to relevant VO concepts. Our approach focuses on directly enriching the existing VIOLIN data nodes (`Vaccine`, `Pathogen`, etc.) with properties and relationships derived from the VO `Resource` nodes. This decision was made to streamline the knowledge graph structure, avoiding the creation of redundant intermediary nodes. By directly attaching relevant semantic information to the original VIOLIN entities, we aimed to simplify querying and improve the efficiency of data retrieval.

Once matched, properties from the `Resource` nodes, such as labels, definitions, and curator notes, were selectively copied and prefixed with “vo_” onto the corresponding VIOLIN data nodes (e.g., `Vaccine` nodes now include properties like `vo_definition`, etc.). This was implemented to enhance query performance and simplify the graph structure. By directly attaching key semantic information from VO to the VIOLIN entities, we avoid the need for complex multi-step traversals through VO nodes during querying, leading to more efficient data retrieval. Furthermore, a `[:VO_REPRESENTATION]` relationship was created from each enriched VIOLIN data node to its matched `Resource` node, establishing a direct link to the VO concept it represents. This relationship provides a clear and explicit connection back to the original ontology term, ensuring traceability and facilitating deeper semantic exploration when needed, while still maintaining a more concise graph structure. This method directly integrates key VO properties into the VIOLIN data nodes, providing semantic enrichment. Finally, to enhance usability, the property keys of the imported `Resource` nodes were updated from the identifiers by Information Artifact Ontology (IAO) to more human-readable labels (e.g., `IAO_0000115` was replaced with `definition`), improving query readability and interpretation of VO concepts within the knowledge graph.

### Node and Relationship Type Definitions

Each node type includes relevant attributes (e.g., `c_vaccine_id`, `c_pathogen_id`, etc.) preserved during the CSV import process. VIOLIN data nodes (`Vaccine`, `Pathogen`, etc.) are further enriched with VO-derived properties (prefixed with “vo_”, such as vo_definition) directly on these nodes. To illustrate relationships within the VIOLIN data, we established connections based on data associations. For example, Vaccines are connected to Pathogens through a `:TARGETS` relationship.

### VaxChat Integration

The integration of Langchain^10^ and Neo4j forms the technological backbone of VaxChat’s core functionality. Langchain efficiently manages the routing of prompts to GPT-3.5 Turbo and processes its outputs, creating a streamlined pipeline for complex language model tasks. Simultaneously, Neo4j provides robust support for query execution and database session management, enabling VaxChat to drive queries and extract relevant data from VaxKG effectively. Together, these technologies create a comprehensive RAG pipeline that translates natural language questions into structured knowledge graph queries and transforms the retrieved data into user-friendly responses. The front-end interface, built using React, emphasizes transparency in the question-answering process. Users can examine both the automatically generated Cypher queries and the raw data retrieved from VaxKG, allowing for verification and a deeper understanding of the information sources underlying each response. The complete codebase for VaxChat is available as an open-source project on the GitHub repository^11^.

## Results

### VaxKG Integration Pipeline

Figure 1 summarizes this overall workflow, illustrating how VIOLIN and VO data sources are transformed, integrated into Neo4j, and subsequently leveraged for semantic queries. We first begin by extracting CSV files from VIOLIN, import the VO, and then unify them in the Neo4j-based VaxKG, which can be queried. Two main use cases are illustrated: (1) performing direct semantic queries in VaxKG, and (2) using VaxChat (GPT-based) to generate Cypher queries and return user-friendly answers from the knowledge graph. While in VaxChat, it’s able to query VaxKG with plaintext user queries by converting a user query into Cypher query. The generated query will then retrieve data from VaxKG. With the retrieved data, another LLM process occurs to generate the answer.

**Figure 1.**
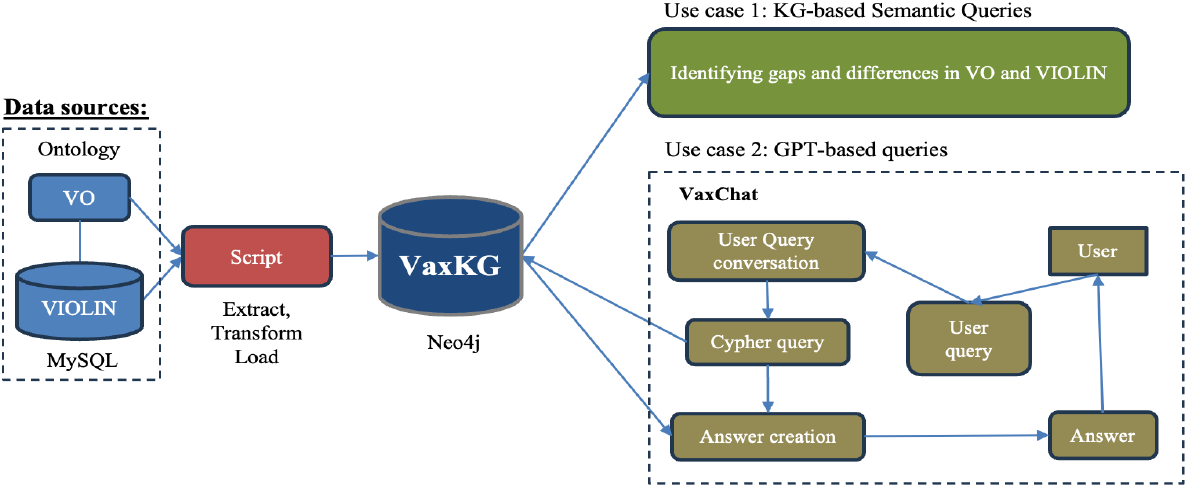
Workflow for constructing and applying the VaxKG knowledge graph.

**Figure 2.**
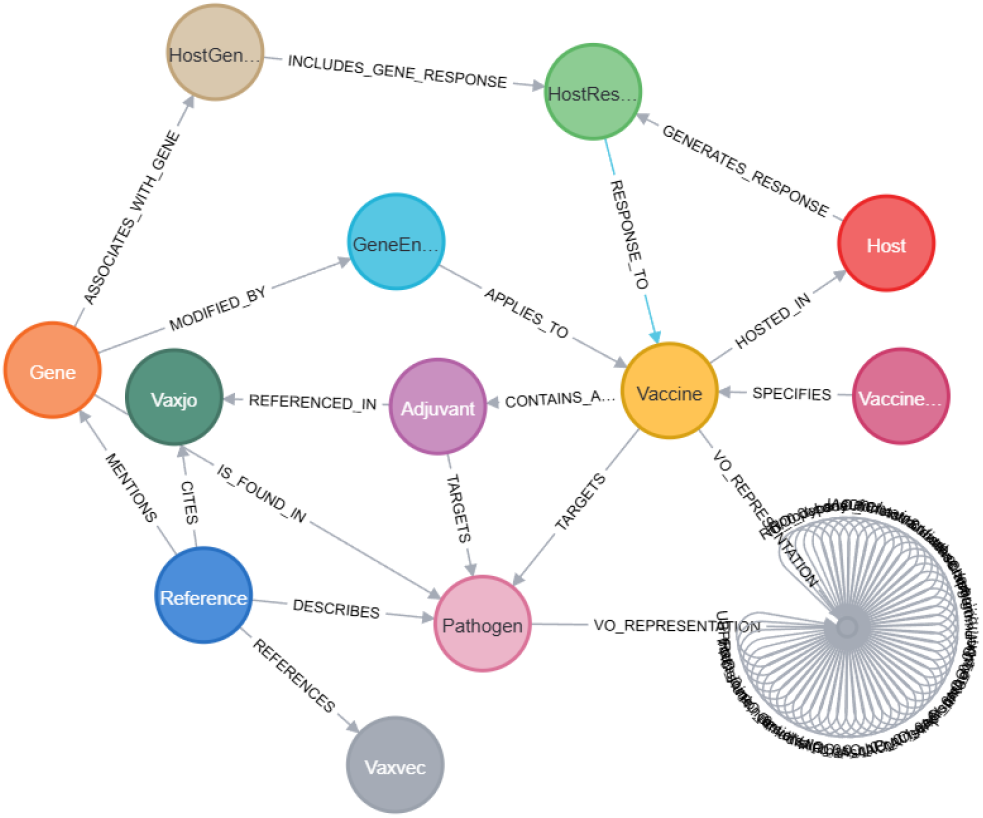
VaxKG Neo4j schema. VaxKG includes 12 core node types and relationships derived from VIOLIN, VO, and the relations between them.

### Knowledge Graph Construction and Composition

Our integrated knowledge graph (KG) was built from 12 core VIOLIN tables with the enrichment from VO. The resulting KG comprises a rich network of interconnected nodes and relationships, significantly enhancing the original VIOLIN database. Nodes in the KG represent diverse entities such as vaccines, genes, and pathogens, while edges capture various relationships such as `INSTANCE_OF`, `INDUCES` and `TARGETS`. Overall, the KG includes 28,123 original VIOLIN data nodes, representing entities such as `Vaccine`, `Gene`, `Pathogen`, and `Host`. It also includes 101,282 `Resource` nodes, imported directly from VO, representing VO classes and properties forming the VaxKG. In terms of relationship count, VaxKG includes 34,453 relationships are generated from the VIOLIN database scheme (e.g., `TARGETS`, `CONTAINS_ADJUVANT`), and 383,511 relationships inherent to the VO itself defining the structure and vocabularies of VO within the VaxKG. There are also 4,335 relationships generated between VIOLIN and VO, creating a total of 412,865 connections altogether within 71 unique relations.

### Use Cases

To demonstrate the utility of integrating the VIOLIN dataset with the VO within a Neo4j knowledge graph, we first performed a series of queries focusing on vaccines against *Brucella* (a bacterial pathogen) and Influenza virus. These queries leveraged the hierarchical structure of VO to enrich and categorize vaccine information extracted from VIOLIN. We later demonstrate how users can navigate through the graph interface to find their targets. Finally, we demonstrate how one can utilize the VaxKG as our knowledge base for our chatbot prototype called VaxChat, allowing the users to uncover comprehensive vaccine information through natural language.

### Semantic Queries: Vaccine-related Queries using Knowledge from VO and VIOLIN

The first query was to investigate the classification of vaccines based on the VO’s hierarchical structure. By identifying vaccines in VIOLIN targeting “*Brucella*” and “Influenza”, and linking them to their corresponding VO representations, we explored the `subClassOf` relationships within VO to categorize these vaccines. The query retrieved a list of *Brucella* and influenza vaccines from VIOLIN, categorized the amount of relevant VO terms.

As shown in Table 1, the VO-only analysis provided a broader view of the conceptual landscape of *Brucella* vaccines, Influenza vaccine, protective antigens, Virmugens and Vaximmutors as defined within the VO. Virmugens are those genes whose mutations would result in pathogen mutants that can be used as live attenuated vaccines^12^. Vaximmutors are vaccine-induced immune factors^13^. For *Brucella* vaccines, we discovered that the ontological architecture of VO covers a slightly wider array of vaccine entities than VIOLIN, identifying 67 distinct vaccines compared to VIOLIN’s 60. This slight difference was caused by VO’s broader hierarchical classification system, which categorizes vaccines based on theoretical relationships and developmental stages, but the two resources still achieved comparable coverage of vaccine candidates. On the other hand, when we examined protective antigens—the molecular components that actually stimulate protective immunity—VIOLIN’s database proved more comprehensive, documenting 29 protective antigens compared to VO’s 19. The inverse relationship of these two reflects VIOLIN’s emphasis on experimental validation and immunological mechanism documentation rather than VO’s comprehensive taxonomic classification. Despite the different approaches, both resources captured a similar number of Virmugens (17 in VO versus 15 in VIOLIN), indicating convergent identification of these important vaccine components. However, the complete absence of Vaximmutor annotations in VO (0 compared to VIOLIN’s 14) reveals a significant gap in VO’s characterization of this particular class of vaccine-related entities for *Brucella*, suggesting fundamental differences in how these resources define and catalog immune response modulators.

**Table 1.** Comprehensive comparative analysis of *Brucella* and Influenza vaccine resources.

|  | *Brucella* |  |  | *Influenza* |  |  |
| --- | --- | --- | --- | --- | --- | --- |
|  | VO | VIOLIN | VO + VIOLIN | VO | VIOLIN | VO + VIOLIN |
| Vaccine Number | 67 | 60 | 59 | 1695 | 412 | 184 |
| Protective Antigen | 19 | 29 | 16 | 51 | 53 | 14 |
| Virmugen | 17 | 15 | 15 | 2 | 1 | 1 |
| Vaximmutor | 0 | 14 | 0 | 0 | 227 | 0 |

The contrast between these resources became even more pronounced when examining influenza vaccines. VO’s ontological framework documented 1,695 influenza vaccine entities, vastly outnumbering VIOLIN’s 412. This substantial difference reflects not only VO’s strength in representing the complex landscape of influenza vaccines by production methods and formulation differences, but also its comprehensive classification of vaccines across diverse host species, including human seasonal influenza, avian influenza, swine influenza, and other host-specific variants. Each host-specific influenza type has its own distinctive vaccine formulations, with VO cataloging these variations as separate entities within its hierarchical structure. Our analysis revealed that VO documented 51 protective antigens for influenza that is comparable to VIOLIN’s 53, suggesting that for extensively studied pathogens like influenza, VO’s coverage of immunological components approaches that of VIOLIN’s specialization. There is, however, another disparity shown in influenza Vaximmutor as VIOLIN documented 227 and VO documented 0. This highlights the fundamental difference in how these resources annotate vaccine-induced immune responses, with VIOLIN providing extensive documentation of experimentally validated vaccine-specific immune modulators that’s absent from VO’s ontological framework. This complementarity shows the value of integrating VIOLIN and VO, as combining their strengths can provide a more comprehensive and structured representation of vaccine data.

Our integrated approach using both VO and VIOLIN resulted in fewer vaccine entities than either resource alone – 59 for *Brucella* and 184 for influenza vaccines. The reduction results from strict matching criteria requiring entities to have corresponding representation in both resources, suggesting that many VO entries lack the experimental validation information like the one in VIOLIN, while some VIOLIN entries lack formal ontological classification like VO. Even more pronounced differences was the reduction in protective antigens in the integrated approach, with only 16 identified for *Brucella* and 14 for influenza—significantly fewer than either resource individually. This suggests potential incompatibilities in how these resources define or identify antigens, with only a subset meeting the criteria for inclusion in both systems.

Despite these reductions, the integration generates meaningful connections between the two different datasets by providing enhanced contextual connections that offer researchers a more nuanced understanding of the vaccine properties. For example, VO’s hierarchical classification of a *Brucella* vaccine can now be linked to VIOLIN’s experimental validation data, allowing researchers to understand both the taxonomic position of a vaccine and its experimentally confirmed properties within a single query. While the integration did not increase the overall number of identified entities in this case, it demonstrates the potential for using VO to provide a semantically enriched context for the information present in VIOLIN by providing classification and functional characteristics of the vaccines. Future work could focus on expanding the integration to potentially uncover additional relationships or insights by leveraging the detailed classifications within VO and refining the matching criteria to address the current limitations in entity mapping between these complementary resources.

### Knowledge Graph Networks Navigation

One of the key benefits of the Neo4j graph database environment lies in its intuitive visual interface, which significantly enhances the exploration and understanding of interconnected data within the knowledge graph, as demonstrated in Figure 3. Panel (b) illustrates the initial output of a Cypher query from panel (a) that is designed to identify vaccines associated with live attenuated resources from the VO. This query reveals a set of matching nodes that results in 189 Vaccine nodes and 111 VO nodes, notably including the *Brucella* RB51 vaccine from the VIOLIN database and a corresponding Resource node from the VO, immediately providing researchers with entities relevant to live attenuated vaccines. The inclusion of the asterisk (`*`) in the `[:subClassOf*]` relationship pattern is crucial. This denotes a variable-length path, allowing the query to traverse any number of ‘subClassOf’ relationships. This powerful feature ensures a comprehensive retrieval of resources that are direct or indirect subclasses of “live attenuated” within the VO hierarchy, thus identifying a broad range of potentially relevant vaccines.

**Figure 3.**
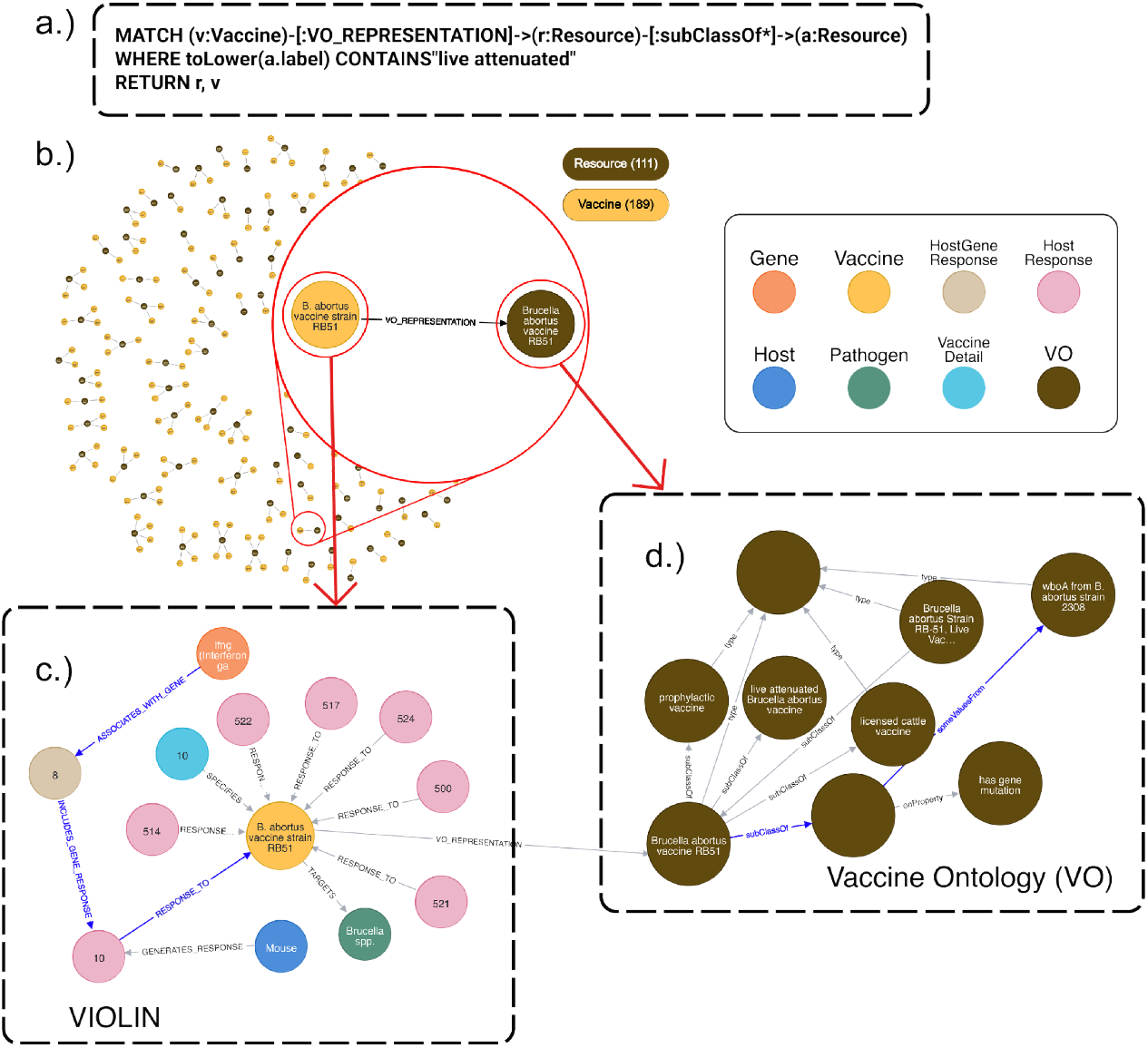
Visual Exploration of the VaxKG in Neo4j. **(a)** A representative Cypher query that queries live attenuated vaccines from VIOLIN and VO. **(b)** 189 live attenuated vaccines were identified from VO and VIOLIN. **(c)** Expansion of the RB51 vaccine node, revealing connected entities within the VIOLIN database and the tracing of a host response node to the IFNg. **(d)** Expansion of a VO Resource node, demonstrating its connection to the *wboA* gene from *Brucella abortus* strain 2308, highlighting a link to the RB51 vaccine.

Progressing to panel (c), we observe the result of expanding the *Brucella* RB51 vaccine node. This expansion unveils the immediate network of related entities within the VIOLIN data, such as host responses. Further expansion of a host response and host gene response node allows for the tracing of connections through blue arrows to specific genes, in this instance, the IFNg gene for IGNg protein (Interferon gamma), offering insights into the host’s reaction to this particular vaccine. While in panel (d), it further showcases the navigational capabilities by illustrating the expansion of a node originating from the VO. This expansion, which includes traversing a ‘subClassOf’ relationship, reveals its connection through blue arrows to the *Brucella abortus wboA* gene that is mutated in the RB51 vaccine strain, highlighting a specific genetic component related to the RB51 vaccine. This visual exploration within Neo4j empowers researchers to intuitively navigate the knowledge graph, discover specific connections between vaccine information in VIOLIN and relevant concepts within the VO, such as the relationship between the *Brucella* RB51 vaccine and the *wboA* gene from a specific bacterial strain. This help fosters a deeper understanding of the complex relationships and pathways within vaccine data and its alignment with standardized ontology terms.

### VaxChat Integration

VaxChat serves as a conversational interface that leverages VaxKG through RAG to provide accessible and accurate vaccine information to users. This system enables researchers to query the knowledge graph using natural language rather than specialized query languages, democratizing access to complex vaccine data.

The architecture of VaxChat follows a prompt engineering approach illustrated in Figure 4. The system employs a two-stage process that first transforms natural language queries into appropriate Cypher queries for Neo4j, and then generates comprehensive answers based on the retrieved data. The prompt engineering involves carefully designed instruction sections that accurately guide the language model in performing these tasks.

**Figure 4.**
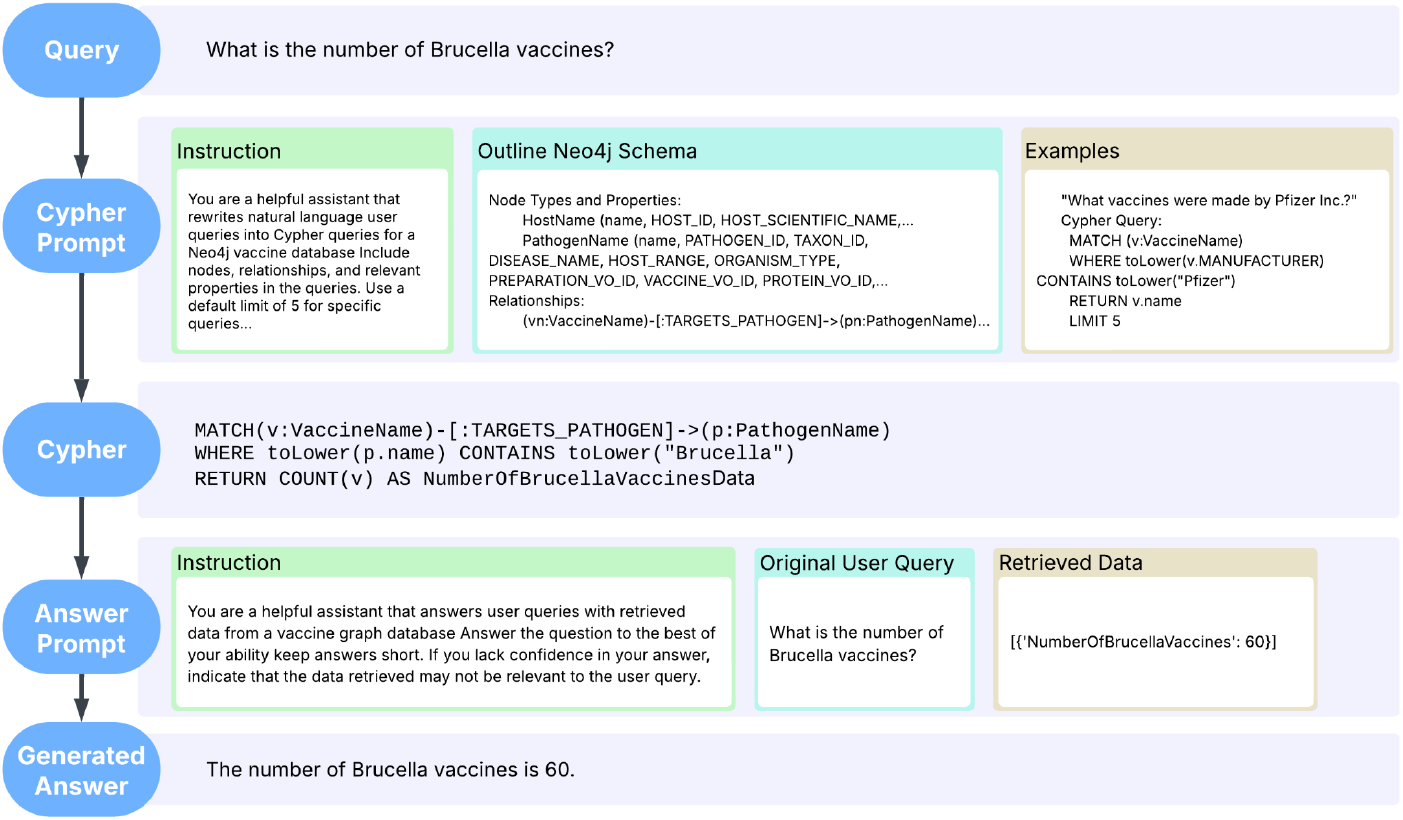
The process of answering a user’s natural language query about the number of *Brucella* vaccines. The query is translated into a Cypher query via a prompt, executed against a Neo4j database, and the retrieved data is used to generate the final answer.

For the Cypher generation component, the language model is provided with detailed context about VaxKG’s structure, including node types, relationships, and property nomenclature. This context enables accurate translation of natural language questions into formal Cypher queries that extract relevant information from the knowledge graph. The prompt includes exemplar pairs of natural language questions and corresponding Cypher translations, establishing patterns for the model to follow when processing new queries.

The answer generation component on the other hand, receives both the original user query and the structured data retrieved from VaxKG. This approach creates the foundation for the model’s responses in factual information from the knowledge graph, significantly reducing the chance of hallucination. VaxChat also maintains transparency by allowing users to inspect both the generated Cypher queries and the raw data retrieved from VaxKG, enabling verification of the information sources underlying each response.

The current implementation of VaxChat utilizes ChatGPT 3.5 Turbo as its underlying language model. However, the system’s modular design allows straightforward substitution with alternative models, including open-source alternatives such as DeepSeek or Llama, providing flexibility for future development and deployment scenarios.

### Evaluation of VaxChat Performance

To provide a structured evaluation of VaxChat’s capabilities, we conducted an assessment based on a set of representative competency questions. This approach tests the system’s ability to answer plausible queries from a vaccine researcher, covering simple data retrieval, aggregation, and handling of linguistic variations such as synonyms and abbreviations. The final natural language answer from VaxChat was manually verified against the VaxKG and categorized.

The results, detailed in Table 2, demonstrate a strong foundational performance but also highlight key areas for improvement in the RAG pipeline. The system achieved a correct or partially correct response on 3 out of the 5 queries. VaxChat excelled at direct questions and aggregation, accurately retrieving specific data points and performing correct counts on the dataset. However, the evaluation revealed a significant weakness in handling linguistic variations. The system was unable to map the common synonym “cows” to the formal term “cattle” or expand the abbreviation “tb” to “tuberculosis,” resulting in query failures. This indicates that while the core knowledge graph is robust, future iterations could leverage a dense entity retrieval model^14^ to better interpret user intent. Such an approach would allow the system to identify and map entities with similar meanings (e.g., “cows” and “cattle”, or “tb” and “tuberculosis”) at the vector level, overcoming the current limitations in handling a wider range of natural language expressions.

**Table 2.** VaxChat Performance on Representative User Queries.

| Question | Generated Answer | Analysis |
| --- | --- | --- |
| How many vaccines target *Brucella*? | There are 60 vaccines that target *Brucella*. | VaxChat successfully executed an aggregation (COUNT) query, accurately matching the underlying data from the VIOLIN database. |
| What is a vaccine that targets *Brucella* for cows | I'm sorry, but the retrieved data does not provide information on a specific vaccine that targets *Brucella* for cows. | The RAG system failed to map the common synonym "cows" to the formal term "cattle" used in the Vaccine Ontology, leading to a failed query. |
| What is a vaccine that targets *Brucella* for cattle | The vaccine that targets *Brucella* for cattle is the "*Brucella abortus* S19 vjbR mutant vaccine." | When the formal term "cattle" was used, the system successfully queried the graph and retrieved a correct example of a licensed cattle vaccine. |
| Is there a vaccine for tb? | The retrieved data does not include a vaccine specifically for tuberculosis (TB). | The RAG pipeline could not expand the common abbreviation "tb" to "tuberculosis". |
| What vaccine targets tuberculosis? | The vaccine that targets tuberculosis is the BCG Vaccine. | The system correctly identified the disease using its full name and retrieved the corresponding vaccine from the knowledge graph. |

## Discussion

This study achieved several transformations towards the integration of vaccine related databases by semantically enriching the database by incorporating VO into VIOLIN 12 core tables. Our approach integrates VO properties into VIOLIN data nodes, creating semantically enhanced entities like Vaccine and Pathogen node types, and establishes `[:VO_REPRESENTATION]` relationships to maintain explicit links to VO concepts. This results in a KG that provides a scalable knowledge base for vaccine research. Furthermore, we have demonstrated the practical usage of this KG through the development of VaxChat, a prototype chatbot application. VaxChat, powered by our KG, offers a more user-friendly natural language interface for accessing complex vaccine information, showing a potential use of a KG that democratizes access to specialized scientific knowledge.

### Advantage of Knowledge Graph Approach

One of the main advantages for using the KG approach is that it enhances the query flexibility and expressiveness. Unlike rigid SQL queries against relational databases, Cypher queries enable the exploration of more complex, and multi-dimensional questions with ease and efficiency. The interconnected structure allows researchers to explore a more complex relationship or pattern within the graph to identify potential novel hypotheses, which would be difficult through the tabular structure of relational databases. Our use case at exploring antigens associated with flu vaccines demonstrate this advantage. Another advantage would be that the integration of VO unlocks semantic reasoning and inference capabilities. The KG architecture lays the foundation for future expansion of the knowledge base to incorporate formal reasoning mechanisms, which unlike relational databases, lacks inherent semantic understanding of the knowledge base^15^.

Furthermore, the VaxKG serves as a crucial knowledge base for the RAG framework similar to how it’s employed by VaxChat. VaxKG’s structured and semantically rich nature allows for the efficient retrieval of relevant information in response to user queries. When a user poses a question to LLM, the RAG process first queries the VaxKG to fetch pertinent nodes and relationships that provide the necessary context. This retrieved knowledge is then used to generate a more accurate and focused answer specifically tailored to vaccine-related topics. By leveraging the VaxKG as its information source, VaxChat avoids relying solely on its pre-trained general knowledge, which might be less specific or even inaccurate for specialized vaccine information. The direct linkages between VIOLIN data and VO concepts within the KG ensure that VaxChat can access and utilize a comprehensive and semantically consistent understanding of vaccine entities and their relationships, ultimately enhancing the quality and reliability of the information it provides to users.

### Limitations

Despite the strength of the current approach to KG generation, it still has some limitations. The quality and completeness of the original VIOLIN database will inherently influence the data quality for VaxKG. Data gaps or inconsistencies within VIOLIN, such as the observed lack of `c_vo_id` values for a subset of vaccines, can limit the extent of ontology mapping and semantic enrichment. Similarly, the VO itself, while comprehensive, may not yet encompass all nuances of vaccine knowledge or may require further expansion in specific areas to fully represent the complexities of the vaccine domain. As the VaxKG grows with the integration of additional data sources and ontology expansions, scalability challenges may arise when there are over million nodes^16^. So, strategies for database optimization and graph processing will be crucial to maintain query efficiency and responsiveness as VaxKG scales. Finally, the process of semantic mapping, while carefully implemented, is inherently complex, therefore, ensuring the accuracy and consistency of semantic interpretations requires ongoing refinement and validation of the mapping process. This work would be even more challenging as the database grows in the future. Future work should focus on addressing these limitations to further enhance the robustness and utility of VaxKG.

### Future Work and Improvement

Future work will focus on several key areas to expand and enhance the KG. A primary direction is the integration of additional relevant datasets beyond the core VIOLIN database. This includes incorporating databases such as clinical trial data, comprehensive drug databases, and insights derived from literature mining efforts. Expanding the ontology coverage by using Immunology Ontology^17^ or Gene Ontology^18^ would also be important to further enrich the VaxKG’s semantic scope. To unlock deeper knowledge discovery, we also plan to apply advanced graph algorithms and analytics to the KG, including exploring graph embedding techniques for machine learning applications, utilizing community detection algorithms to uncover hidden relationships within the vaccine knowledge network. We recognize the importance of user accessibility, so future efforts will be directed towards developing interactive visualization dashboards and user interfaces to facilitate a user-friendly data exploration user interface that allows for easier knowledge discovery for a wider range of researchers. This work includes enhancement to VaxChat by incorporating more advanced natural language processing and KG navigation capabilities, further allowing researchers for easier access to their necessary vaccine data and information.

A key focus of our future work is the multi-faceted, rigorous evaluation of VaxChat’s accuracy. While the initial competency questions provide a baseline, a more formal approach is needed. First, we will establish a rigorous gold standard benchmark by collaborating with domain experts to create and validate a comprehensive set of complex questions and their correct answers. This benchmark will specifically target the types of ambiguity and incompleteness identified in our initial analysis. Second, we will perform a deep error analysis on VaxChat’s incorrect responses. This involves dissecting the RAG pipeline to determine the point of failure: was it an error in the initial Cypher query generation, or a misinterpretation by the language model during final answer synthesis? The insights from this analysis will directly guide targeted improvements in our prompt engineering and data mapping strategies. Finally, we will conduct a comparative evaluation, benchmarking VaxChat’s accuracy and task-completion time against traditional methods, such as a researcher using the VIOLIN website directly. This will provide quantitative evidence of the system’s utility in accelerating knowledge discovery. The findings from our accuracy analysis will drive improvements to VaxChat’s natural language processing and KG navigation capabilities, creating a more robust, reliable, and user-friendly tool for the vaccine research community.

## Acknowledgements

The author(s) declare financial support was received for the research, authorship, and/or publication of this article. This work has been supported by the NIH-NIAID grants U24AI171008 and the Undergraduate Research Opportunity Program (UROP) at the University of Michigan.

